# Effect of land use, habitat suitability, and hurricanes on the population connectivity of an endemic insular bat

**DOI:** 10.1101/2020.12.18.423522

**Authors:** Camilo A. Calderón-Acevedo, Armando Rodríguez-Durán, J. Angel Soto-Centeno

## Abstract

Habitat loss and fragmentation are a leading cause of vertebrate population declines and extinction. Urbanization and natural disasters disrupt landscape connectivity, effectively isolating populations and increasing the risk of local extirpation particularly in island systems. Puerto Rico, one of the most isolated islands in the Caribbean, is home to 13 bat species that have been differentially affected by disturbance during the Anthropocene. We used circuit theory to model the landscape connectivity within Puerto Rico with the goal of understanding how fragmentation affects corridors among forested areas. Models combined species occurrences, land use, habitat suitability, and vegetation cover data to examine connectivity in the endemic bat *Stenoderma rufum*, and also at the bat community level across the island. Urbanization in Puerto Rico affected bat connectivity overall from east to west and underscored protected and rustic areas for the maintenance of forest corridors. Suitable habitat provided a reliable measure of connectivity among potential movement corridors that connected more isolated areas. We found that intense hurricanes can disrupt forest integrity and affect connectivity of suitable habitat. Some of the largest protected areas in the east of Puerto Rico are at an increasing risk of becoming disconnected from more continuous forest patches. The disruption of corridors that maintain connectivity on the island could explain previous findings of the slow post-hurricane population recovery of *S. rufum*. Given the increasing rate of urbanization, this pattern could also apply to other vertebrates not analyzed in this study. Our findings show the importance of maintaining forest integrity, emphasizing the considerable conservation value of rustic areas for the preservation of local biodiversity.

## 1. Introduction

Untangling the processes that affect biodiversity is a fundamental endeavor in ecology and conservation biology. Due to the fast pace of global change, processes like habitat conversion or fragmentation during the Anthropocene have become increasingly important as potential drivers of biodiversity loss in terrestrial ecosystems (Ceballos & Ehrlich 2002; Meyer et al. 2016; Torres-Romero et al. 2020). Island systems are in special need for biodiversity conservation assessments because many islands are considered biodiversity hotspots due to their high number of endemic species (Mittermeier et al. 2011). Yet, a significant proportion of native insular biodiversity is often threatened by an ever-growing human footprint that has led to a significant loss of habitat (Schoener et al. 2004; Barnosky et al. 2011; Pimm et al. 2014; Turvey et al. 2017; Turvey & Crees 2019; Andermann et al. 2020). Terrestrial vertebrates on islands in the Caribbean are further threatened by an increased frequency in extreme climatic events such as hurricanes, which can have short- and long-term impacts on their populations (Donihue et al. 2020). Therefore, evaluating the events that may lead to population changes in island systems is key to assessing how biological communities are structured across the landscape and examine their responses to anthropogenic and natural disturbances. Understanding how the remaining habitat connects across the landscape provides an exceptional opportunity to address issues of habitat conversion and fragmentation to examine how potential biological corridors may aid in conservation efforts.

The Caribbean island of Puerto Rico, itself a small archipelago of some 140 small islands and cays, contains seven different types of forest habitats (Gannon et al. 2005; Miller & Lugo 2009). These forests are subject to various levels of anthropogenic and natural pressures across the island derived from urban development concentrated in the northeast and south-central areas (Guzmán-Colón et al. 2020), and agricultural development located primarily in the west and southwest. Across Puerto Rico, there are 118 protected and specially protected rustic areas with an extension of over 3300 km^2^. These areas are classified as commonwealth forests, nature reserves, federal reserves, and non-governmental protected areas (Gould et al. 2008a; Gould 2009; Gould et al. 2011). Combined, the protected and rustic areas form corridors that span from east to west and connect all forest types in the main island of Puerto Rico. Rustic areas lack conservation management but enclose most of the government protected areas and act as habitat buffer zones. Thus, rustic areas help separate State Forests by providing isolation from disturbance (i.e., urbanized areas or main roads), and due to their agricultural, archeological, ecological, or natural resource value are somewhat protected from development and urbanization (Gould et al. 2008a; Junta de Planificación 2015; Gould et al. 2017). The east coast of Puerto Rico includes El Yunque National Forest, the largest protected area of the island. El Yunque National Forest and the nearby Carite State Forest (Fig. 1 A and B, respectively) contain subtropical wet and rain forests and lower montane wet and rain forests. Notwithstanding, these large protected areas are under anthropogenic pressure and are affected by urbanization (Guzmán-Colón et al. 2020). On the center and west of the island, the Toro Negro State Forest (Fig. 1 C) connects to the Maricao State Forest (Fig. 1 D) and the Río Abajo State Forest (Fig 1. E). These central and western localities encompass subtropical wet and lower montane wet and rain forests.

**Figure 1.**
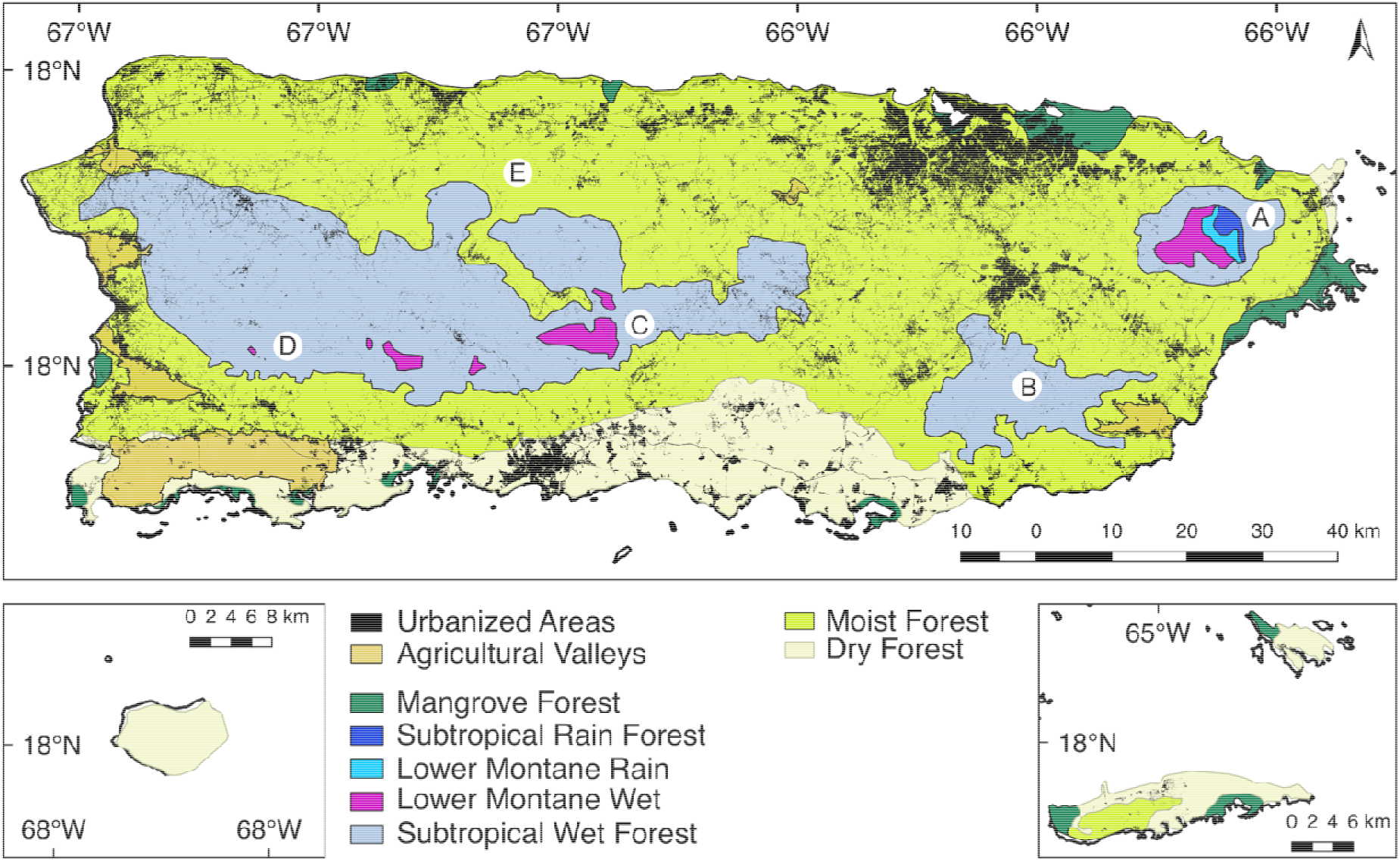
Forest types, agricultural valleys and urbanized areas of Puerto Rico. A) El Yunque National Park protects 5 different kinds of forests and many smaller ecosystems; B) Carite State Forest; C) Toro Negro State Forest; D) Maricao State Forest and adjacent areas; E) Río Abajo State Forest and adjacent areas.

The local fauna of the island is affected by anthropogenic disturbance, but also endure the seasonal pass of hurricanes and tropical storms, which may lead to population declines and potentially an increased risk of extirpation. In Puerto Rico, hurricanes have caused the decline or the potential extirpation of some bat species (Gannon & Willig 1994, 1998, 2009; Jones et al. 2001; Rodríguez-Durán & Vazquez 2001; Rodríguez-Durán et al. 2020). These effects have been observed in the most abundant and generalist species, *Artibeus jamaicensis*, in and the endemic and far less abundant, *Stenoderma rufum*, which populations took nearly 5 years to recover to pre-hurricane levels following hurricane Hugo in 1989 (Gannon & Willig 1994, 1998). *Stenoderma rufum* is known from the Virgin Islands where it is considered rare (Gannon et al. 2005; Kwiecinski & Coles 2007). In Puerto Rico, this species primarily roosts in trees and occurs in forests throughout the island (Genoways & Baker 1972; Gannon et al. 2005). Drastic declines of this species in Puerto Rico could potentially affect populations in the smaller Virgin Islands by reducing the likelihood of post-hurricane recovery via recolonization from this source population. The negative effects of hurricanes on *S. rufum* in Puerto Rico could also be exacerbated by the poor continuity of forest patches connecting local demes between the east and west of the island.

We aimed to model the population connectivity of *Stenoderma rufum* on Puerto Rico to examine the potential effects of anthropogenic (i.e. forest fragmentation) and natural (i.e. hurricanes) pressures. Given the roosting ecology, low population density, and slow post-hurricane recovery of *S. rufum*, we hypothesized that this species is adapted to dense forests that can be disrupted by anthropogenic or natural disturbance. Herein, we provide 1) an inference of the potential areas that offer the highest connectivity, 2) documentation of possible corridors with suitable habitats that connect protected areas, and 3) examination of the potential effect of hurricanes on the connectivity of *S. rufum* across Puerto Rico. Our population connectivity modeling approach sheds light into the importance of different forested areas for future conservation efforts of this insular endemic species and potentially other vertebrates.

## 2. Materials and Methods

We modeled the landscape connectivity of the red fig eating bat (*Stenoderma rufum*), an endemic species on Puerto Rico, based on occurrence localities across the island. Our area of interest spans the main island of Puerto Rico, covering all seven types of forest and areas under any form of protection. Focusing on *S. rufum* in Puerto Rico was advantageous because this island has the largest, best monitored, and most continuous populations of the species compared to satellite populations in the Virgin Islands (Gannon & Willig 1998, 2009; Rodríguez-Durán 1998; Kwiecinski & Coles 2007; Rodríguez-Durán & Padilla-Rodríguez 2010; Rodríguez-Durán & Feliciano-Robles 2016; Gómez-Ruiz & Lacher 2017; Rodríguez-Durán et al. 2020). Additionally, land use categorization and forest type distribution in Puerto Rico are well known and the Puerto Rico Gap Analysis Project has available datasets of high resolution (Gould et al. 2008a; Gould 2009). Our landscape modeling approach used electrical circuit theory implemented in the software Circuitscape v4.05 to examine the landscape connectivity of *S. rufum* under different scenarios (Shah & McRae 2008; McRae et al. 2008). Circuit theory calculates the flow of current between pairs of nodes (i.e. species localities) in the landscape, the latter being a continuous surface with resistance or conductance values that indicate different potential pathways (i.e. habitat corridors) for animals to move through (McRae et al. 2008). By calculating the flow of current across each node one can identify areas of interest that can potentially help maintain the population connectivity of a species across the landscape (Carroll et al. 2012; Theobald et al. 2012; Dutta et al. 2016; Mallory & Boyce 2019; Osipova et al. 2019).

### 2.1 Study Area and Data Sources

The main island of Puerto Rico has an area of 8,948 km^2^ of which 46% is dedicated to urban and barren areas, and agricultural valleys and pastures; forests, woodlands and shrublands account for 54% of the total area (Gould, 2009). To the east, the island is under pressure by rapid urban development and an increasing human footprint (Guzmán-Colón et al. 2020). Although the protected and rustic areas offer connectivity between the main forests through the island (Fig. 1).

We obtained all known georeferenced localities of *Stenoderma rufum* (N = 46) in Puerto Rico, including places of capture and roosts from the published literature and GBIF records (Gannon et al. 2005; Rodríguez-Durán & Feliciano-Robles 2016; GBIF.org 2020). For the connectivity analyses, we used land use and protected areas data from the Puerto Rico Gap Analysis (Gould et al. 2008b, 2008a; Martinuzzi et al. 2008; Gould et al. 2011). Also, publicly available climate and elevation data grids from WorldClim v2 were used as environmental variables to estimate the ecological niche of *S. rufum* (Fick & Hijmans 2017). Finally, three Normalized Difference Vegetation Index (NDVI) layers representing vegetation cover information from January 2017, October 2017, and January 2020 were used to examine the effects of forest cover change on connectivity (Vermote & NOAA CDR Program 2019). All spatial grid data was scaled to a resolution of 1 km.

### 2.2 Models of Population Connectivity

We modeled the connectivity of 26 unique localities of *S. rufum* (Fig. 2, Supplementary Table 1) under 4 scenarios, 1) connectivity based on a resistance layer using different categories of land use; 2) connectivity using the protected areas as a conductance layer; 3) connectivity of *S. rufum* based on a conductance layer that accounted for habitat suitability estimated using ecological niche modeling (ENM); and 4) connectivity based on a layer of conductance using NDVI across three time snapshots that reflected forest cover change following a recent natural disturbance (i.e., hurricanes Irma and Maria) and its effect on the landscape connectivity.

**Figure 2.**
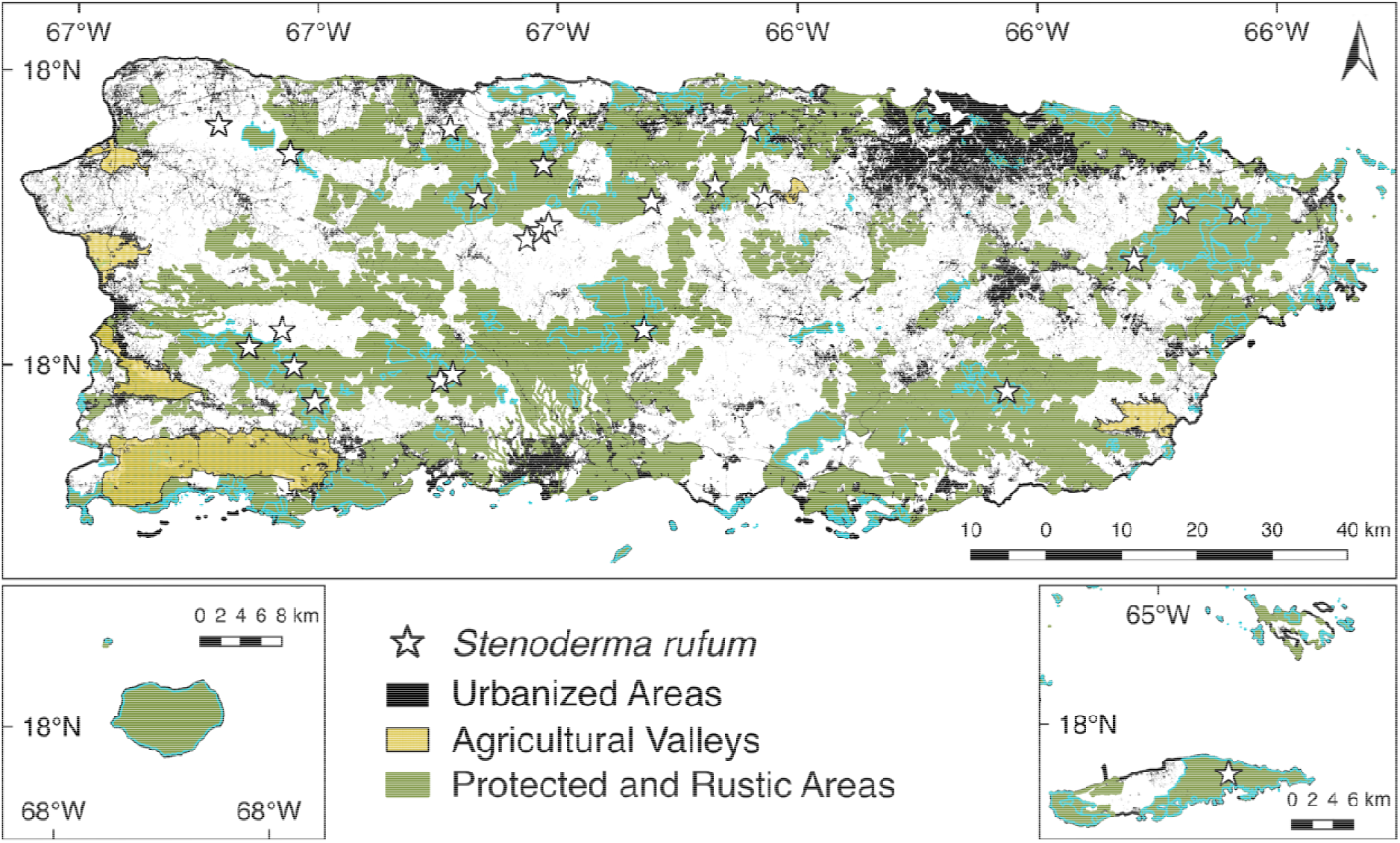
Record localities (i.e., stars) of *Stenoderma rufum* used to produce connectivity models in Puerto Rico. Map shows urbanized areas, agricultural valleys, and protected and specially protected rustic areas that were used for calculating functional connectivity. Light blue contours denote the area with government protection following Gould (2009).

#### 2.2.1 Effects of land use on bat connectivity

To understand the effect of land use on bat connectivity, we used the land use classification grid of the Puerto Rico Gap Analysis Project (Fig. 2; Gould et al. 2008b; Martinuzzi et al. 2008; Gould 2009). We reclassified cell values from low to high resistance using a value of 1 for any forest, rural and non-urbanized area, which would provide the least resistance to bat movement in the environment. We also set a resistance of 3 for agricultural valleys and 5 for highly urbanized areas, defined as any built up and non-vegetated areas resulting from human activity based on Gould et al. (2008b), which usually correspond to large urban areas and show the highest resistance. This approach allowed us to understand how resistant the landscape is to bat mobility. We calculated the connectivity also in terms of the landscape being a conductive surface in which protected and rustic areas provide higher conductance than developed areas. Here, grid values were classified as 0 for urban areas and agricultural valleys (i.e., low conductance), a value of 1 for rustic areas because they share a direct edge with developed and urbanized areas (i.e., intermediate conductance), and a value of 2 for government protected areas (i.e., high conductance).

We also wanted to examine whether land use helps predict congruent patterns of connectivity at the community level for bats in Puerto Rico. For this, we used 137 unique localities (Supplementary Table 2) for the 13 bat species known from the island, which represent 3676 records from GBIF, and calculated connectivity as described above.

#### 2.2.2 Land connectivity accounting for habitat suitability

To examine the connectivity based on the suitable habitat required by *S. rufum*, we first estimated habitat suitability using an ecological niche modeling (ENM) approach under a maximum entropy framework. Specifically, we modeled the potential distribution of *S. rufum* to use as input for a conductance model. ENM was generated using MaxEnt v3.4.1 (Phillips et al. 2006; Phillips & Dudík 2008) and included 19 current climate variables and elevation available in WorldClim 2 (Fick & Hijmans 2017). We used a regularization multiplier value of 0.5 and let the algorithm of Maxent converge using the “auto features” setting, which resulted in the selection of linear, quadratic, and hinge (LQH) as the final features used for modeling. Species observations were randomly partitioned into 75% training and 25% testing sets using 100 bootstrap replicates. The resulting average habitat suitability map produced from the ENM of *S. rufum* was used as a conductance grid. We calculated this suitable habitat connectivity between all pairs of the 26 unique localities of *S. rufum*.

#### 2.2.3 Effects of natural disturbance on bat connectivity

We used NDVI data from three specific time periods, January 2017, October 2017 and January 2020 (Vermote & NOAA CDR Program 2019) to create three independent conductance models. These three models were used to compare the connectivity among localities of *S. rufum* before, immediately after, and 26 months after the most recent large scale storms that affected Puerto Rico in September 2017 (i.e., hurricanes Irma and Maria). We used the data range of NDVI values in our grid as conductance values for Circuitscape, with higher NDVI values representing better conductance across the landscape.

## 3. Results

Our results from the different connectivity analyses of *Stenoderma rufum* showed four important areas that promote connectivity among roosting localities across the island (Fig. 3; refer to Fig. 1 for name places). The three analyses suggest limited connectivity between the eastern localities of El Yunque National Forest and Carite State Forest. Furthermore, both of these eastern localities show low connectivity with the central and western localities, i.e., Río Abajo and Toro Negro State Forests and Maricao State Forest, respectively.

**Figure 3.**
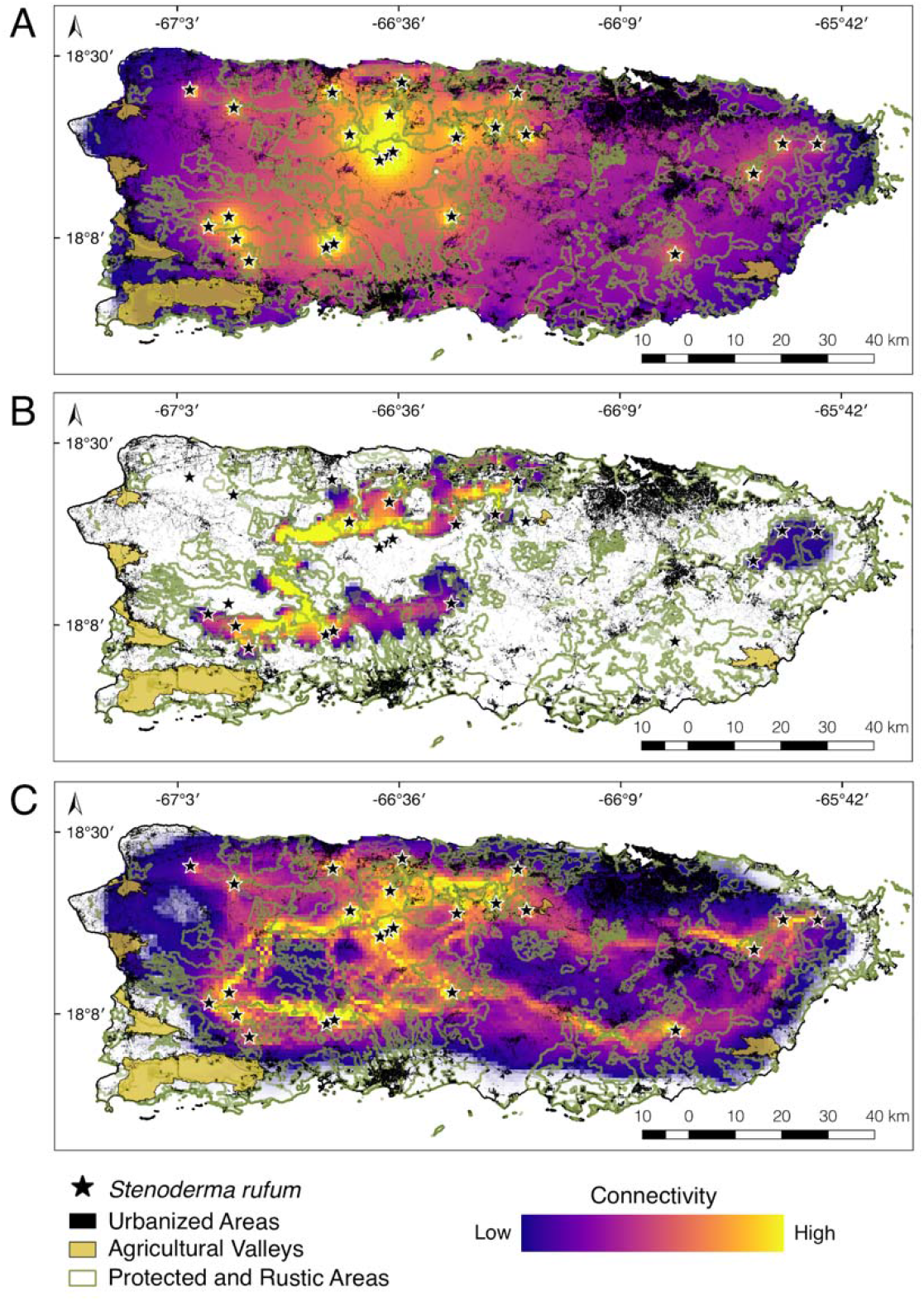
Landscape connectivity of *Stenoderma rufum* across the island of Puerto Rico estimated in Circuitscape. Three models were developed using A) resistance from land use data, B) conductance mediated by protected areas, and C) habitat suitability derived from an ecological niche model. Warmer colors indicate higher levels of connectivity, black and white areas denote no connectivity.

When land use alone was included as a resistance layer, i.e. differential permeability in the landscape depending on the type of land use, the model produced an even connectivity gradient across the island (Fig. 3A). This gradient generally showed lower connectivity in the coastal areas that gradually increased towards inland forested areas. As predicted, only the urbanized, developed, and agricultural areas showed gaps in connectivity among *S. rufum* localities, and increased connectivity was observed among localities that cluster close together. Connectivity was concentrated primarily in the west of Puerto Rico, where urbanization is lower than in the eastern areas (Fig. 3A).

We observed a similar pattern to the one described above when using protected and rustic areas as conductance surfaces to model the connectivity of *S. rufum*. Localities within protected forests in the west of Puerto Rico were well connected from north to south but disconnected from the scattered localities in the east (Fig. 3B). Despite the expansiveness of protected and rustic areas in the east including El Yunque National Forest and Carite State Forest, these lacked intermediate suitable forests that can act as ecological corridors to connect with the central and western regions of the island. This result was mirrored when analyzing the whole bat community of Puerto Rico. Areas in the west of the island serve as corridors that connected habitat latitudinally, while El Yunque National Forest remained disconnected (Supplementary Figure S1).

The ecological niche model of *S. rufum* revealed that suitable habitat for this species concentrates in subtropical moist forest, particularly throughout lower montane, rain and moist forests (Supplementary Fig. S2). Habitat suitability decreased rapidly over coastal areas, lowlands, agricultural valleys, and subtropical dry forests. When habitat suitability was used as a conductance layer, two main corridors were observed via the north and south of the island that link El Yunque National Forest with the western forests stretching to Toro Negro, Río Abajo, and Maricao State Forests (Fig. 3C). We found more north to south connections between the State Forests to the western part of the island, whereas El Yunque National Forest connected to the nearby Carite State Forest only by a narrow strip of suitable habitat running parallel to the southeastern coast (Fig. 3C). The western forests (i.e., Maricao, Río Abajo, and Toro Negro) connected north to south creating a loop of forests with multiple narrow corridors (Fig. 3C). This loop overlaps with the connectivity corridor created by protected and rustic areas (Fig. 3B).

Connectivity analyses also reflected the potential negative effects of hurricanes Irma and Maria on *S. rufum* across the landscape (Fig. 4). Estimates of connectivity using pre-hurricane NDVI data (i.e., January 2017) as a conductance grid showed an even connectivity gradient throughout the island (Fig. 4A). This pattern was congruent with our expectations based on models produced using land use data. There was a decrease in vegetation cover immediately after the hurricanes (i.e., October 2017; Liu et al. 2018). This loss of vegetation cover observed in October 2017 disrupted connectivity and created gaps from north to south among Toro Negro, Río Abajo, and Maricao forests in the west (Fig. 3B). Models produced using NDVI data 14 months post-hurricane (i.e., January 2020) showed that forest connectivity increased nearly to pre-hurricane levels.

**Figure 4.**
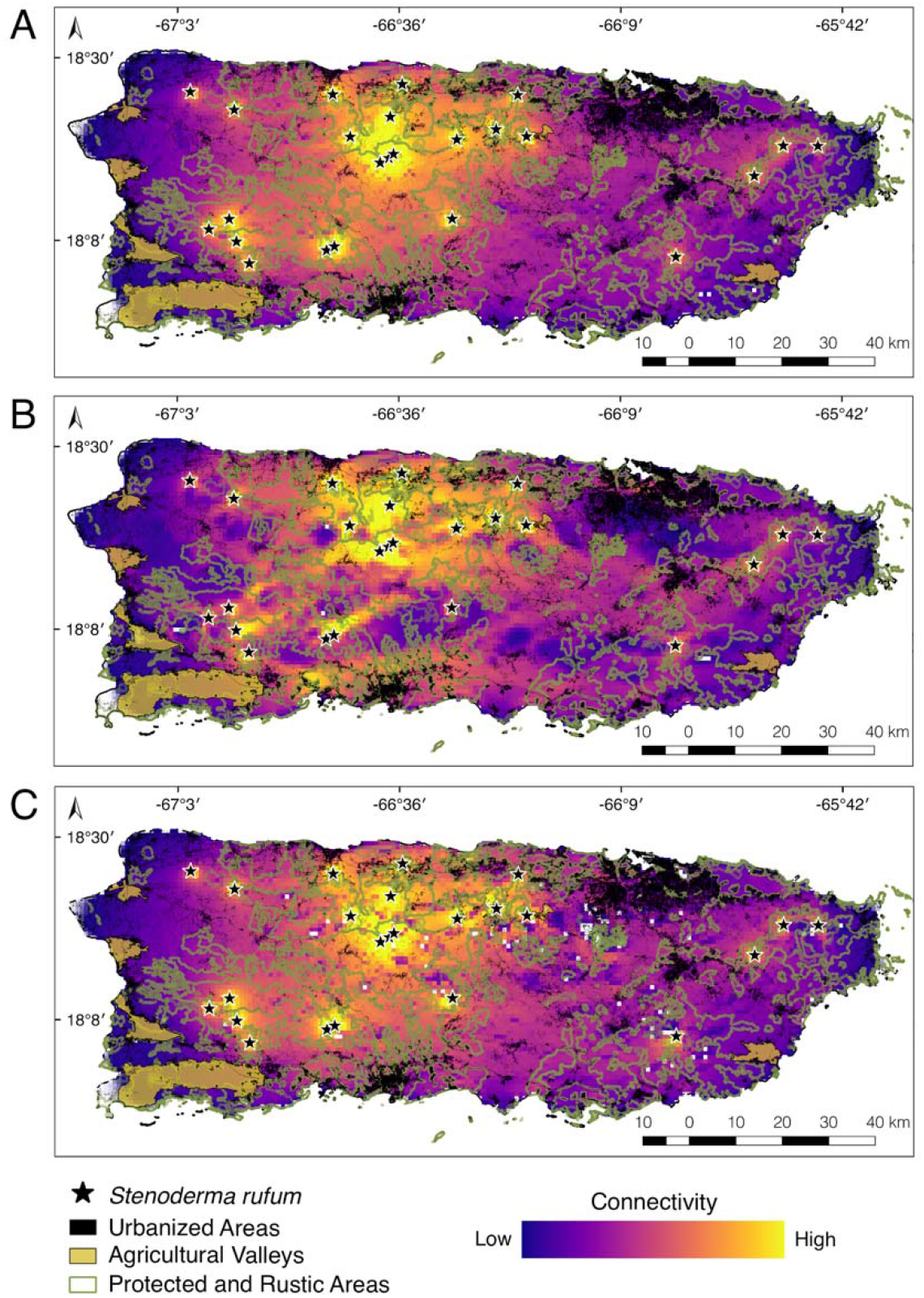
Connectivity of *Stenoderma rufum* based on conductance calculated using the Normalized Difference Vegetation index values from A) January 2017, B) October 2017 and C) January 2020 to reflect availability of vegetation cover before and after hurricanes Irma and Maria.

## 4. Discussion

The current biodiversity crisis stems from several causes, among them, habitat fragmentation is one of the main drivers of biodiversity loss in continental and insular mammal fauna (Ceballos & Ehrlich 2002). Habitat fragment size and isolation within the landscape are important factors for quantifying the response of bat populations to habitat loss. Urbanization and loss of vegetation cover either via anthropogenic or natural causes may affect species in different ways (Meyer et al. 2016). For example, while specialist and/or rare species could be displaced, common species may flourish or recover easily, effectively modifying the composition of insular bat communities. Although the current population trend of *S. rufum* is unknown (Rodríguez-Durán 2016), our results suggest that land use, natural disasters, and the availability of suitable habitat remaining play a role in maintaining the connectivity across the landscape for this bat. The maintenance of connectivity in *S. rufum* is important because of its solitary habits, ephemeral roosting preferences, and small home range of ca. 2 ha (Gannon 1991; Gannon & Willig 1994). Until 30 years ago, *S. rufum* was known in Puerto Rico from only two localities. The rareness of this species and its propensity to become extirpated from forests where it was once abundant make it imperative to better understand how *S. rufum* maintains its resilience in the face of habitat loss.

The model that takes into account the available suitable habitat of *S. rufum* irrespective of land use (Fig. 3C) presents the most likely forest corridors for the species because it shows connectivity based on abiotic requirements. Conversely, the models constructed using land use data and vegetation cover (Fig. 3A and Fig. 4A-C), should be taken as a best case scenario of connectivity because they assume that the species could move across any habitat in the landscape. Even though these models represent gradients of connectivity throughout the island, they do not account for suitable habitat. Our results nevertheless highlight the importance of the western protected and specially protected rustic areas in Puerto Rico for providing connectivity corridors between the north and south coasts. Furthermore, these findings emphasize the need for the preservation of the suitable habitat of *S. rufum* to ensure the connectivity among local demes. The importance of these habitats in Puerto Rico extends beyond this species to the whole bat community of the island. Under the different modeled scenarios, the protected areas in the west always remain connected, which may emphasize the importance of underdeveloped or low urbanized areas to maintain corridors for bat population connectivity (Fig. 2 and Fig. 3). The only decrease in connectivity in this region was observed when models included vegetation cover data immediately after a natural event (Fig. 4; September 2017). The influence of a high intensity category-4 storm, hurricane Maria, underscores the potentially detrimental effects that loss of vegetation cover can have on bat populations.

Several important points can be gleaned from the modeling scenarios we presented. First, habitat on the east of Puerto Rico could be highly susceptible to future changes in land use and urbanization because of the already fragmented nature of the landscape. Second, the interconnecting corridors between the preserved forests to the east and west can be vital for maintaining gene flow in *S. rufum* and may help in species recovery after natural disturbances. Third, the western forests (i.e., Maricao, Río Abajo, and Toro Negro) form a well-connected network of habitat and each may serve as a hub or refugium for the long term persistence of populations. The landscape connectivity based on land use data show similar patterns when examining *S. rufum* and also for the entire bat community of Puerto Rico. This is particularly important in the center of the island, where the corridor narrows (Fig. 3B and Supplementary Figure S1). If these protected areas maintain ecological connectivity, they could represent important movement corridors for local fauna.

Recently, Guzmán-Colón et al. (2020) proposed a connectivity model using only protected areas and forest patches representing the home ranges of several vertebrates in Puerto Rico. Their findings are highly congruent with our results, despite that they were produced using different methods. Guzmán-Colón et al. (2020) found that the human footprint in several scattered patches outside of the protected areas is “very low” and indicative of habitat fragmentation. Furthermore, they documented a network hub in the central northwest of Puerto Rico. Although the analysis of Guzmán-Colón et al. (2020) focused on other vertebrates that are potentially less vagile than bats, their results support our conclusions that the western forests provide a local hub for *S. rufum* connecting with other areas of the island and that the eastern localities could be under the pressure of habitat fragmentation and loss of connectivity.

In contrast to Guzmán-Colón et al. (2020), we combined data from rustic areas in addition to government protected forests to provide estimates of connectivity. Because of their designation, rustic areas are under less pressure of development in the future. Despite that these rustic areas have a relatively low human footprint, they could still be used for agricultural purposes in the short term. We believe that rustic areas are of high importance for vertebrate populations across the island because they encompass a significant amount of suitable habitat that can improve connectivity among local demes across Puerto Rico. These rustic areas, therefore, represent important points for land conservation efforts.

Studies documenting bat responses to habitat fragmentation often show species specific results. On landbridge islands of Lake Gatún, Panama, researchers have documented that abundances of insectivorous Neotropical bats often decline in response to habitat fragmentation, whereas plant visiting bats tend to increase (Meyer & Kalko 2008; Estrada-Villegas et al. 2010). In both cases, previous results suggested that small forest fragments seem to hold significant conservation value and forest integrity was proposed to be of high conservation priority. On Puerto Rico, and other Caribbean islands, forest integrity is disrupted by hurricanes, which pose intense and frequent natural disturbances in addition to ongoing anthropogenic forest fragmentation. These hurricanes cause rapid changes that alter forest integrity and reduce vegetation cover particularly affecting old forest stands more significantly than younger ones (Feng et al. 2020). Although by now mostly recovered, some of the areas most affected by hurricane María in 2017 and having the longest lasting residual effects include El Yunque National Forest, and Carite, Maricao, Río Abajo, and Toro Negro State forests (see Fig. 4 in Feng et al. 2020).

Previously, the bats of Puerto Rico have shown a decline in response to hurricane activity. Bat populations showed a significant reduction in abundance and sampling across habitats revealed a decrease in species richness (Jones et al. 2001). In contrast to the patterns described above based on habitat fragmentation on landbridge islands of Panama, the effects of hurricane Georges in 1998 produced the opposite short-term effect on Puerto Rico, with insectivores increasing in abundance and plant visiting bats showing a decrease. In addition to shifts in species composition, recent studies highlight the susceptibility of bats to rapid disturbance with the possible extirpation of the frugivorous *Artibeus jamaicensis* following hurricane Maria (Rodríguez-Durán et al. 2020). Gannon and Willig (1994) showed that population recovery of *S. rufum* took five years to reach pre-hurricane levels at El Yunque National Forest after hurricane Hugo in 1989. Following hurricane María, *S. rufum* was not present on forest fragment sites surveyed near the eastern metropolitan areas where it had been captured before (Rodríguez-Durán unpub. data). Given the magnitude of forest cover loss and the post-hurricane disruption of habitat corridors (Fig. 3 and Fig. 4), it is likely that the absence and slow recovery of *S. rufum* from eastern Puerto Rico results from the lack of connectivity across the landscape. Maintaining the habitat corridors coupling western forests with the more isolated eastern forests in Puerto Rico could help facilitate the recolonization of *S. rufum* and may be key to maintaining resilience among populations.

Even though bats are the only native mammals remaining on many Caribbean islands (Soto-Centeno & Steadman 2015), our study takes a first step into understanding the potential effects of land use, habitat suitability, and natural disasters to estimate intra island bat population connectivity. One notable finding of these models is that suitable interconnected habitat for *S. rufum* is located primarily on the west of Puerto Rico, despite that all information about this bat has come primarily from the east at El Yunque National Forest (Gannon et al. 2005). It is important to note that our models estimate connectivity only from the perspective of habitat availability and additional data could help clarify the use and importance of habitat corridors on the island. A clear next step for validating our findings is to incorporate landscape genetics to untangle whether the corridors estimated herein help maintain gene flow of *S. rufum* across the island. Deciphering whether populations in each of the main protected forests hold important genetic variability and the directionality of gene flow can shed light into the source-sink dynamics of local demes following habitat disruption by hurricanes. Integrating this approach with connectivity models can help identify areas where populations might shift their ranges as suitable habitat changes under different climate change scenarios (Razgour 2015). Additional studies on *S. rufum* should include a boots-on-the-ground approach targeting the habitat corridors we predicted herein and tracking the movement of individuals across these corridors to confirm their suitability as communing or foraging areas. The models presented offer a new perspective for studying intra island bat communities and highlight the potential value of rustic areas for increasing forest integrity and areas of significant conservation value for *S. rufum*, and potentially other vertebrates on Puerto Rico.

## Acknowledgements

We thank N. de la Sancha and M. Gehara for their advice on connectivity analyses, A. Paz for her help with geographic data processing, and R. D. Barrilito for the motivating discussions on modeling. Work by JASC was partly supported by the National Socio-Environmental Synthesis Center (SESYNC) under funding received from the National Science Foundation DBI-1639145 and by a Rutgers University Research Council Award. Work by CCA was funded by a postdoctoral scholarship at the Soto Lab of Bat Biology (SLaBB) in Rutgers University.

## Supplementary Material

### Supplementary Figures

**Supplementary Figure S1.**
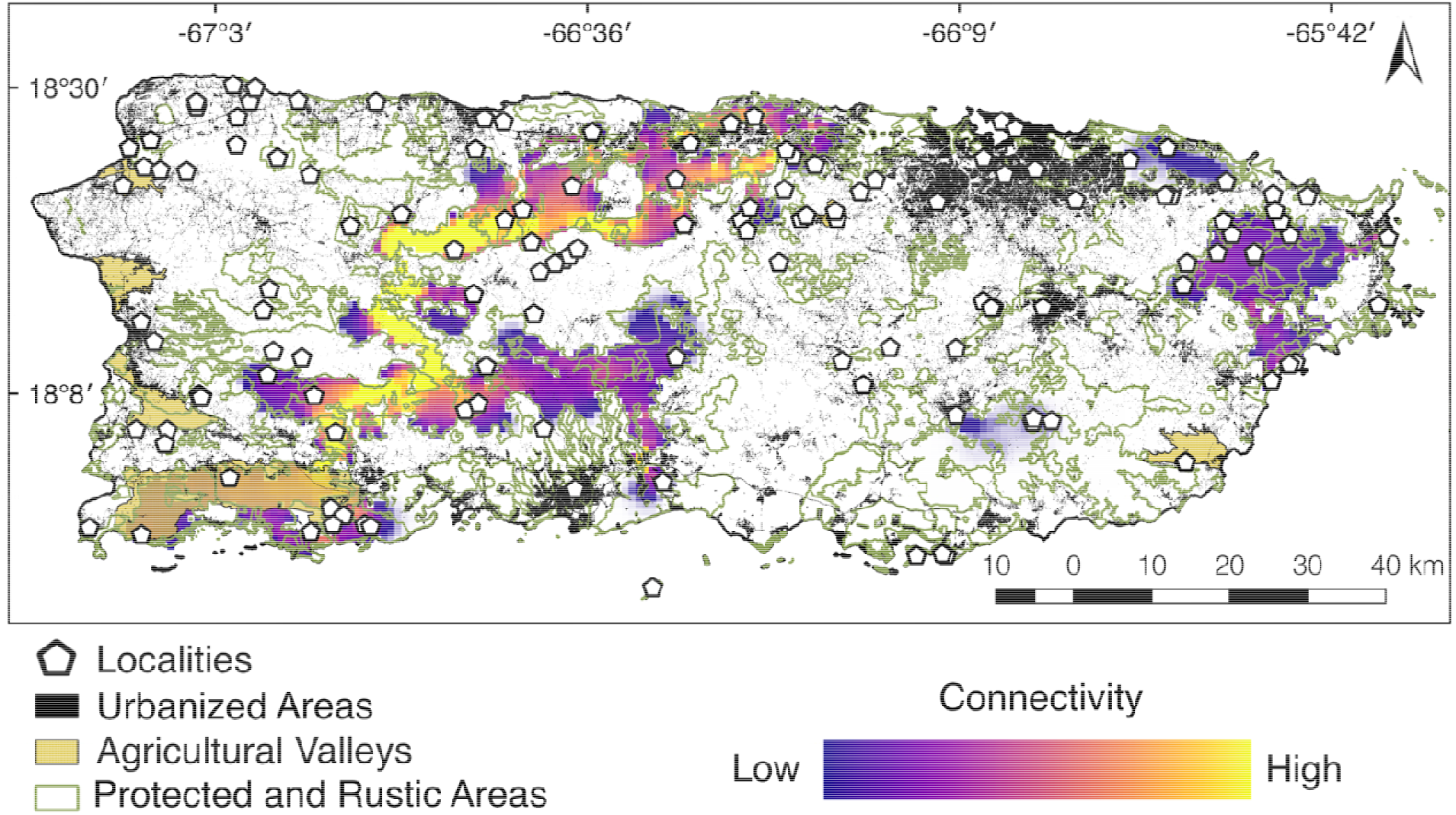
Connectivity models calculated for 137 bat localities in Puerto Rico (Gannon et al. 2005) using protected and especially protected specially protected rustic areas as conductance.

**Supplementary Figure S2.**
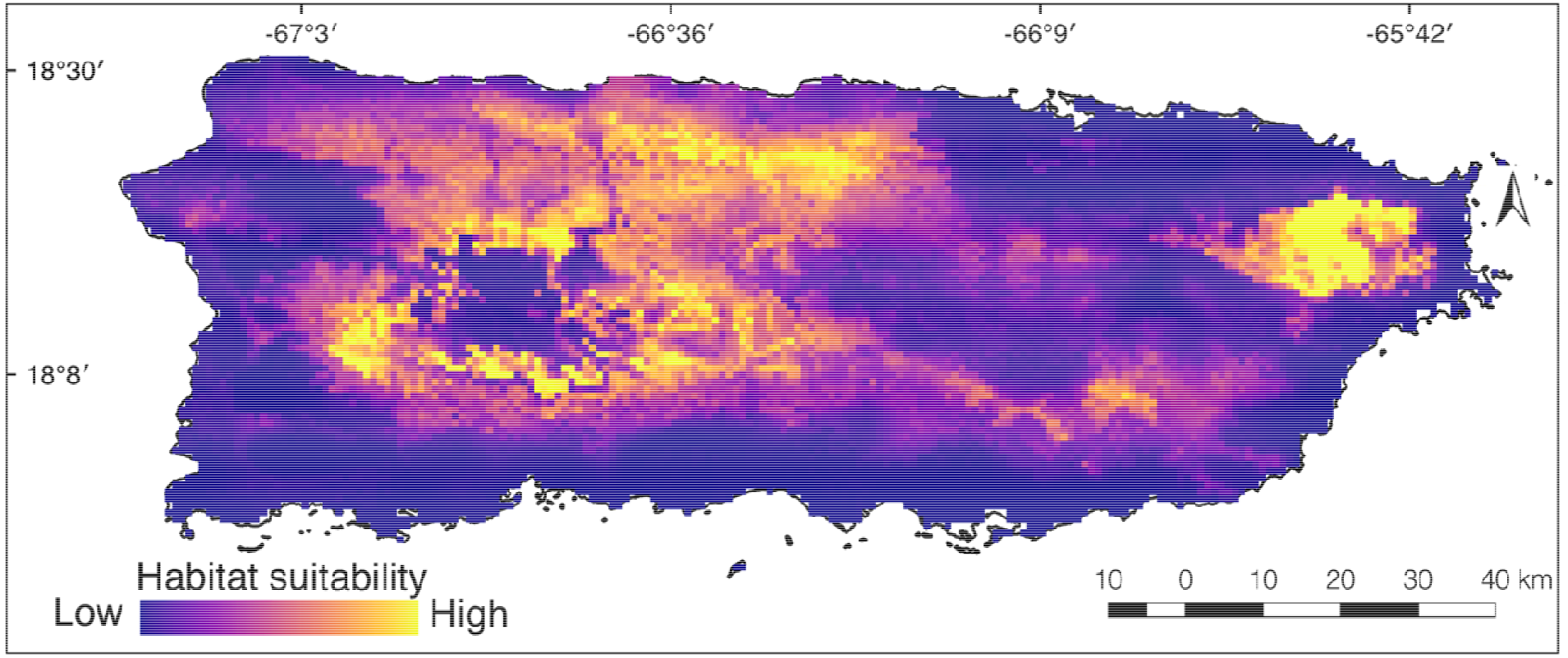
Ecological niche model of *Stenoderma rufum* in Puerto Rico based on 19 WorldClim climate variables summarizing regional temperature and precipitation.

### Supplementary Tables

Supplementary Table 1. Georeferenced localities of *Stenoderma rufum* used in the Environmental Niche Modeling analysis.

Supplementary Table 2. Georeferenced localities of representing the bat community in Puerto Rico, taken from Gannon et al. (2005) and Rodríguez-Durán & Feliciano-Robles (2016).

